# The methylation of *SDC2* and *TFPI2* defined three methylator phenotypes of colorectal cancer

**DOI:** 10.1101/2021.03.03.433833

**Authors:** Ruixue Lei, Yanteng Zhao, Kai Huang, Kangkang Wan, Tingting Li, Haijun Yang, Xianping Lv

## Abstract

Methylation-based noninvasive molecular diagnostics are easy and feasible tools for the early detection of colorectal cancer (CRC). However, many of them have the limitation of low sensitivity with some CRCs detection failed in clinical practice. In this study, the clinical and pathological characteristics, as well as molecular features of three methylator-groups, defined by the promoter methylation status of *SDC2* and *TFPI2*, were investigated in order to improve the performance of CRC detection. The Illumina Infinium 450k Human DNA methylation data and clinical information of CRCs were collected from The Cancer Genome Atlas (TCGA) project and Gene Expression Omnibus (GEO) database. CRC samples were divided into three groups, HH (dual-positive), HL (single positive) and LL (dual-negative) according to the methylation status of *SDC2* and *TFPI2* promoters. Differences in age, tumor location, microsatellite instable status and differentially expressed genes (DEGs) were evaluated among the three groups and these findings were then confirmed in our inner CRC dataset. The combination of methylated *SDC2* and *TFPI2* showed a superior performance of distinguishing CRCs from normal controls than each alone. Samples of HL group were more often originated from left-side CRCs whereas very few of them were from right-side (*P* < 0.05). HH grouped CRCs showed a higher level of microsatellite instability and mutation load than other two groups (mean nonsynonymous mutations for HH/HL/LL: 10.55/3.91/7.02, *P* = 0.0055). All mutations of *BRAF*, one of the five typical CpG island methylator phenotype (CIMP) related genes, were found in HH group (HH/HL/LL: 51/0/0, *P* = 0.018). Also there was a significantly older patient age at the diagnosis in HH group. Gene expression analysis identified 37, 84 and 22 group-specific DEGs for HH, HL and LL, respectively. Functional enrichment analysis suggested that HH specific DEGs were mainly related to the regulation of transcription and other processes, while LL specific DEGs were enriched in the biological processes of extracellular matrix interaction and cell migration. The three defined mathylator groups showed great difference in tumor location, patient age, MSI and ECM biological process, which could facilitate the development of more effective biomarkers for CRC detection.

## Introduction

Colorectal cancer (CRC) is responsible for over 1 million new cases every year and around 700,000 deaths occurred worldwide, making it the third most frequently diagnosed cancer [1,2]. In China, the incidence and mortality of CRC has been witnessed with an increasing trend of 12.8 in 2003 to 16.8 per 100,000 in 2011 and 5.9 in 2003 to 7.8 per 100,000 in 2011, respectively [3]. It is believed that CRCs represent a heterogeneous group of tumors characterized by complex multifactorial phenotypes and no single risk factor is responsible for the developing of CRC [4,5]. Many factors including diet, tobacco smoking, microbial, overweight and obesity, genetic factors, as well as metabolic and other exposures can alter the risk of getting CRC [6–9]. Nearly half of colorectal cancer incidence and mortality was attributable to unhealthy diets such as low vegetable and fruit intake, high red and processed meat intake, and alcohol drinking *etc*., in China in 2012 [8]

Syndecan-2 (*SDC2*), as one of the syndecan family of heparan sulfate proteoglycan, has been demonstrated playing an important role in cancer progression through regulation of cell adhesion, proliferation, and migration in many researches [10–13]. Tissue factor pathway inhibitor-2 (*TFPI-2*), belongs to the Kunitz-type serine proteinase inhibitor family and is thought to be functional in the regulation of extracellular matrix digestion and re-modeling by inhibiting a broad spectrum of serine proteinases [14,15]. Unlike the tumorigenic behaviors of SDC2 in colon cancer cells, *TFPI2* has been shown as a tumor suppressor gene in several malignant tumors [16–19]. However, both promoters of the two genes were found with frequently hyper-methylated status in colon cancer cells compared to normal tissue cells in a few epi-genomics studies [20,21]. The frequently aberrant DNA methylation of SDC2 and *TFPI2* makes them promise biomarkers for the early detection of CRC [22–24] and hence were also used as CRCs diagnostic biomarkers in our preliminary clinical trials. Molecular subtyping analysis based on DNA methylomics data identified a subset of CRCs, CpG island methylator phenotype (CIMP), which is characterized by significant hyper-methylated CpG islands of tumor suppressors [25]. When combined with the clinicopathological parameters and molecular characteristics, CIMP tumors showed significant associations with *BRAF* mutations, MSI-H, female sex, right-sided tumor location, and age [25]. Additionally, prognostic analysis showed that CIMP-high patients presented a worse prognosis than CIMP-low, which suggested CIMP could be a predictor of prognosis of CRC [26].

CRC is a disease with high heterogeneity and many differences were observed among CRCs raised from proximal (right) or distal (left) colon. For example, right-sided colon cancers were reported an increased incidence of proximal migration, while it was inversed for rectosigmoid tumors [27]. What’s more, the incidence rates between proximal colon and distal colon also differ in age and gender [28]. These data reflect an extensively distinct molecular pathogenesis between the cancers originated from these two anatomical locations, which might be related to significant impact on tumorigenesis in these respective sides [28]. In terms of genomic features, proximal carcinomas are characterized by more often microsatellite instable (MSI), frequent *BRAF*-mutated and expressing the CIMP phenotype [29,30]. Therefore, tumor location would be an important factor of biological heterogeneity and it should be reasonable to group CRC into right-sided (proximal) and left sided (distal) ones [31].

Several studies have suggested a better performance of combined multi-targets for CRC early detection than single biomarker [32–34]. However, during clinical practice, some CRC samples was detected with only single or no target positively, reflecting the preference of different targets in distinguishing CRCs from normal samples. In this study, we first classified CRC samples into three methylator groups, *SDC2/TFPI2* double positive group (HH, Hypermethylation-hypermethylation), *SDC2/TFPI2* single positive group (HL, hypermethylation-hypomethylation) and *SDC2/TFPI2* double negative group (LL, hypomethylation-hypomethylation) according to the promoter methylation status of *SDC2* and *TFPI2*, which were previously determined as dual-targets for CRC early detection by our custom window-sliding method (data not published). The clinicalpathological parameters and molecular features were then evaluated by inner and outer samples including TCGA COAD/READ cohorts, GEO datasets as well as our D311 CRC dataset. These findings indicated that it might be reasonable to define three *SDC2/TFPI2* methylator groups according to their methylation status and will benefit the development of more effective methylated biomarkers.

## Methods

### Data preparation

The level 3 methylation data, raw read-count of RNA-seq and clinical information of colon adenocarcinoma and rectum adenocarcinoma patients were retrieved from The Cancer Genome Atlas (TCGA) data portal (https://portal.gdc.cancer.gov/) by using the TCGAbiolinks R package [35]. The platform of methylation data from TCGA is Illumina Human Methylation 450 Beadchip (450K array) and we also searched the Gene Expression Omnibus (GEO; http://www.ncbi.nlm.nih.gov/geo/) for eligible datasets that are generated by 450K array. Two GEO datasets, GSE48684 [36] and GSE79740 [37], were then downloaded because of their available clinical information.

Empirical Bayes (EB) batch adjustment along with two step quantile normalization method [38] was conducted for batch effect removal before GSE48684 and GSE79740 datasets were merged as one set. Missing values of the 450k array were inferred and fulfilled by the Bayesian Network structure learning algorithm [39]. All samples without clinical information were removed and the information of preprocessed data used in this study was presented in **Table 1**.

**Table 1.**
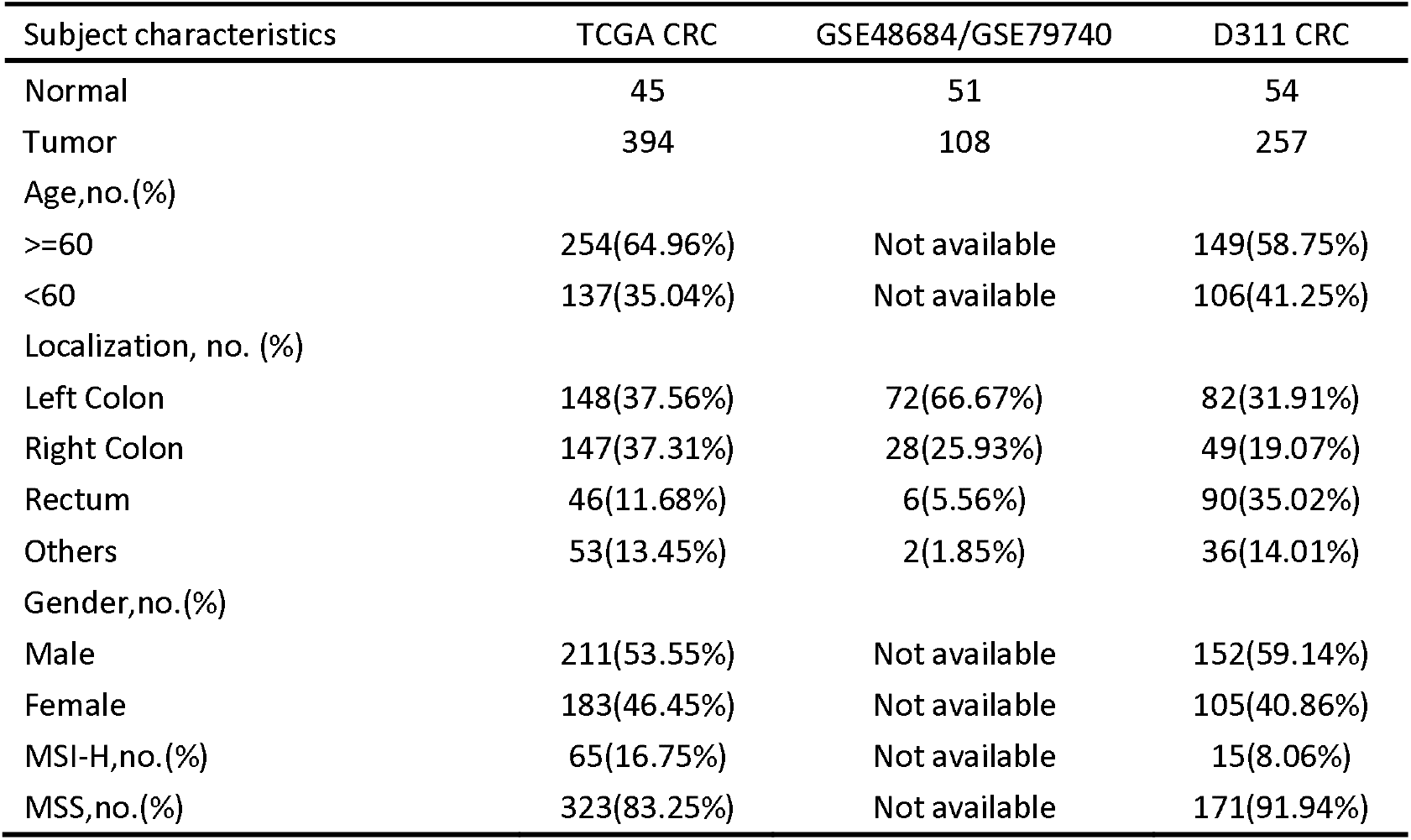
Clinical characteristics of subjects.

### Patient samples

Fresh-frozen colorectal cancer tissues (n=300) and colorectal mucosa (n=55) tissues were collected at Zhongnan Hospital of Wuhan University at the time of surgery. All objects recruited signed a written informed consent and their final diagnosis were determined based on colonoscopy or histological test. Participants who undertook any chemotherapy or radiotherapy, or had incomplete information were excluded. The collected information consist of age, gender, tumor size, tumor location, grade and MSI status. Detailed demographic and clinical features of the subjects were listed in **Table 1**. We defined rectosigmoid, descending colon, and splenic flexure tumors as left-sided cancer, whereas hepatic flexure and ascending colon tumors were grouped as right-sided cancer [40]. This study was approved by the medical ethics committee of Zhongnan Hospital of Wuhan University (No.2019099).

### Methylation-specific PCR experiments

The genomic DNA were extracted by UnigeneDx FFEE DNA extraction kit according to manufacturer’s instruction. Target genes in tumor tissue were captured using previously reported technology with some modification [24]. Tissue derived genomic DNA was chemically modified by sodium bisulfite to convert unmethylated cytosine to uracil while leaving methylated cytosine unchanged. Methylation-specific PCR (MSP) was used to determine the methylation status of *SDC2* and *TFPI2* in normal and tumor tissue DNA, β-actin [41] was used as internal control. Specific primers and probe for the target region of *SDC2* and *TFPI2* was designed as showing in **Table 2**. We use the cycle threshold (Ct) value to determine the methylation status of these two genes and the values for tissue samples were considered “invalid” if the ACTB Ct was greater than 36.00 and methylated *SDC2/TFPI2* were considered “detected” if the Ct values were less than 45.00. For samples with no amplification curve of the MSP occurred after 45 cycles, Ct value was assigned 45.00. Three MSP replicates was done for each sample and the average Ct value was used for further analysis.

**Table 2.**
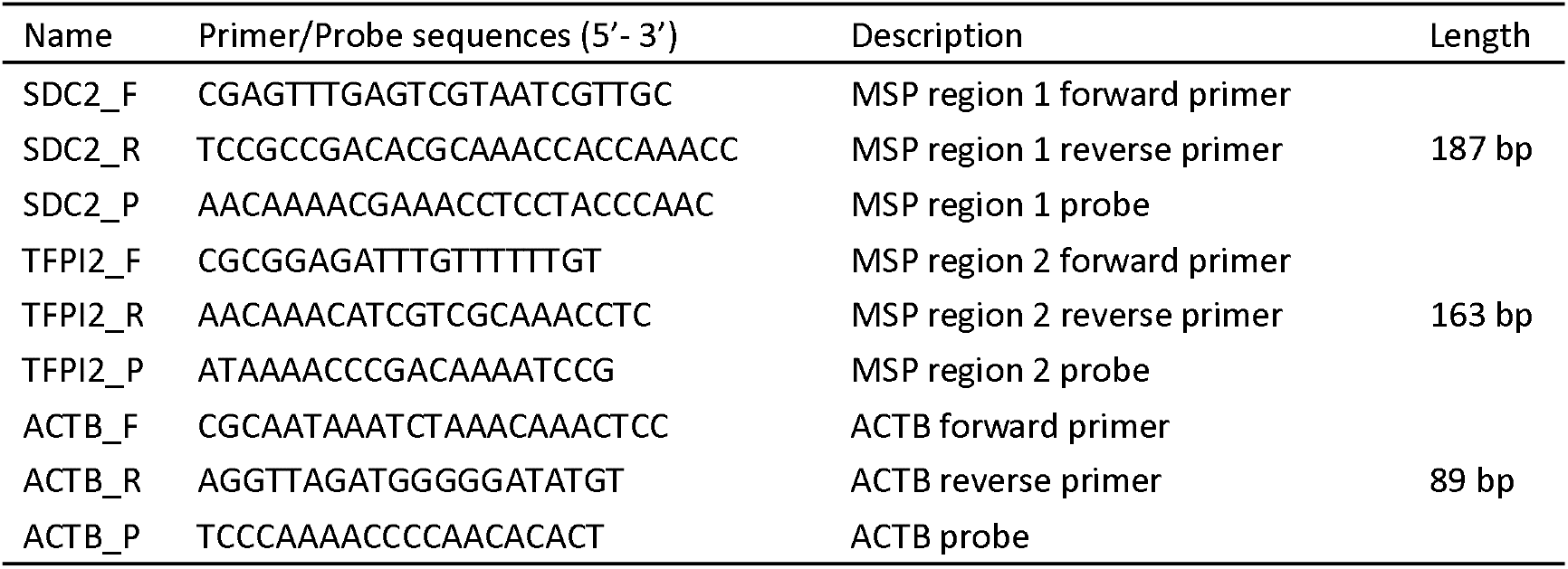
Primers and probes used in this study

### Identification of differentially expressed genes

Level 3 RNA-seq data of TCGA CRCs was preprocessed before differentially expressed analysis by removing low expressed genes whose expressions were zero among more than 90% samples. DESeq2 [42] (V1.30.0) package was used to perform a pairwise comparison between all the three methylator groups for the identification of differentially expressed genes (DEGs). Adjusted P values were calculated by false discovery rate (FDR) method. Genes with adjusted P value < 0.05 were selected as DEGs and used for further functional enrichment analysis. We used GeneCodis [43] for the enrichment analysis of GO biology process (BP) and KEGG pathway for identified DEGS_◦_

### Statistical analysis

Comparisons for two paired or unpaired samples were performed for continuous variables using paired or unpaired student t-test. Kruskal-Wallis nonparametric analysis were used for multi-group comparisons of continuous variables. For categorical variables, fisher’s exact test was applied to determine if there are nonrandom associations between *SDC2/TFPI2* methylator groups and clinical characteristics, such as age, sex and tumor location. For 450k array data, it was proposed to use the thresholds of 0.2 and 0.8 to define hypo- and hyper-methylation [44,45]. In this study, we grouped CpG sites into two categories, hypo-methylated and hyper-methylated based on their β values with the threshold of 0.2. Our D311 CRC samples are defined as detectable negative if Ct values of both the dual-targets are > 38, otherwise they are defined as detectable positive. Sensitivity and specificity were calculated as follows:

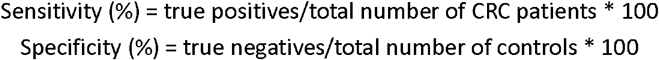

The performance of the diagnostic models were evaluated by the area under receiver operating characteristic (ROC) curve (AUC) and 95% confidence interval (CI). All statistical analyses were performed using R software (version 3.6.0).

## Results

### The methylation of *SDC2* and *TFPI2* showed good discrimination of CRCs from normal controls

Probes within 2 kilo-base upstream of the transcription start site of *SDC2* and *TFPI2* were selected and their mean methylation levels were then calculated according to the β values of 450k array by using TCGA and GEO CRC samples. We only used the probes with Δβ (βTumor-βNormal) >=0.3 and identified 4 and 7 probes in the promoter of *SDC2* and *TFPI2* (**Table 3**). The average β values of these filtered probes was used as the methylation level of *SDC2* and *TFPI2* (herein termed as SDC2_P and TFPI2_P). A lower Δβ value was found for *TFPI2* than *SDC2* which might attribute to its higher background methylation level on normal controls (**Figure 1A**). We also observed a higher sensitivity and lower specificity for *TFPI2* than *SDC2* (**Figure 1B**). When combining *SDC2* and *TFPI2*, the diagnostic sensitivity was improved (**Figure 1B**), which demonstrated that dual-target biomarkers could distinguish CRCs from normal controls better than single target. In our clinical outcomes, CRCs showed significantly higher methylation levels than normal (**Figure 1C**). The pattern of specificity and sensitivity, as well as the combined sensitivity for *SDC2* and *TFPI2*, were also in line with our former results.

**Table 3.**
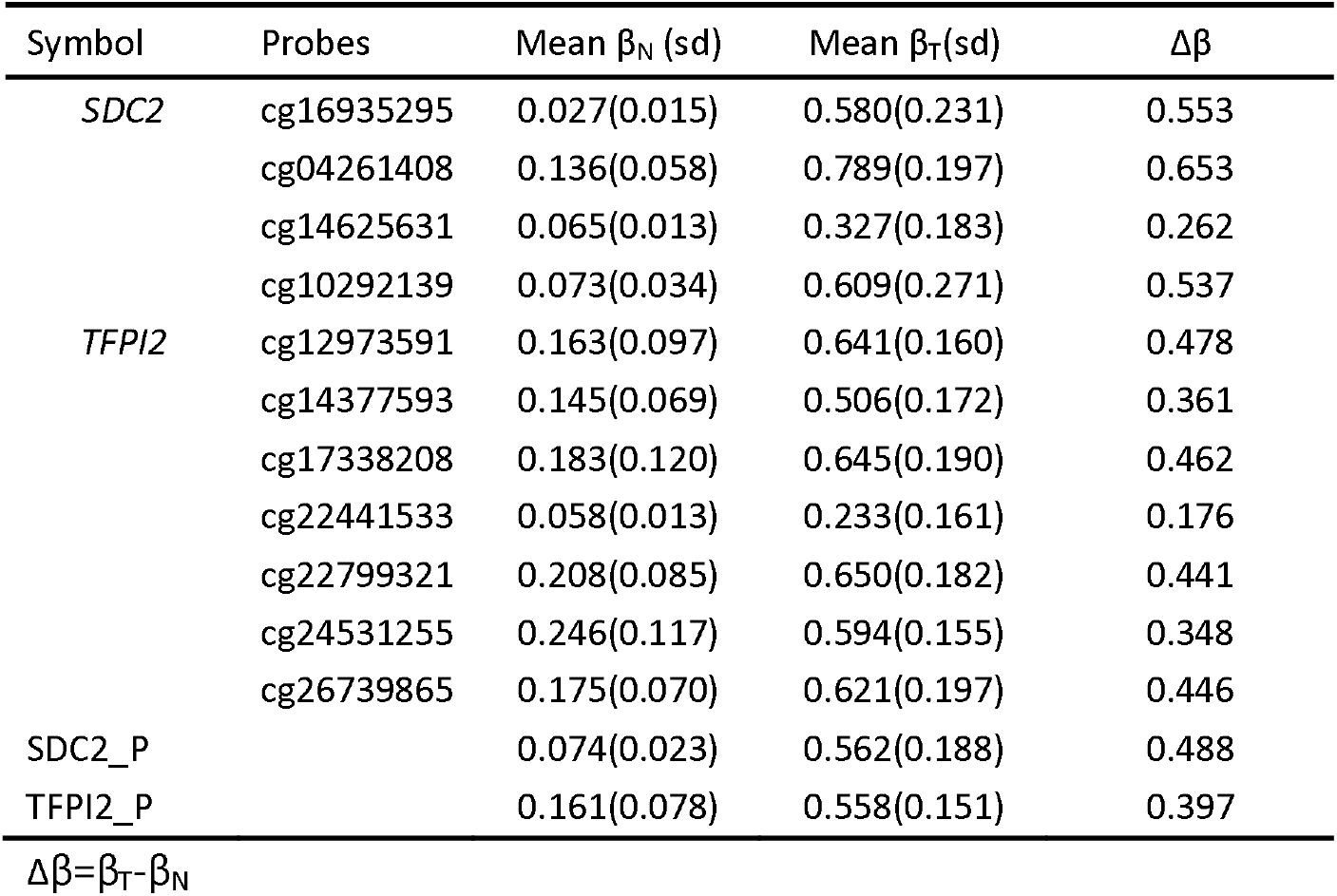
Probes identified in the promoter of SDC2 and *TFPI2*.

**Figure 1.**
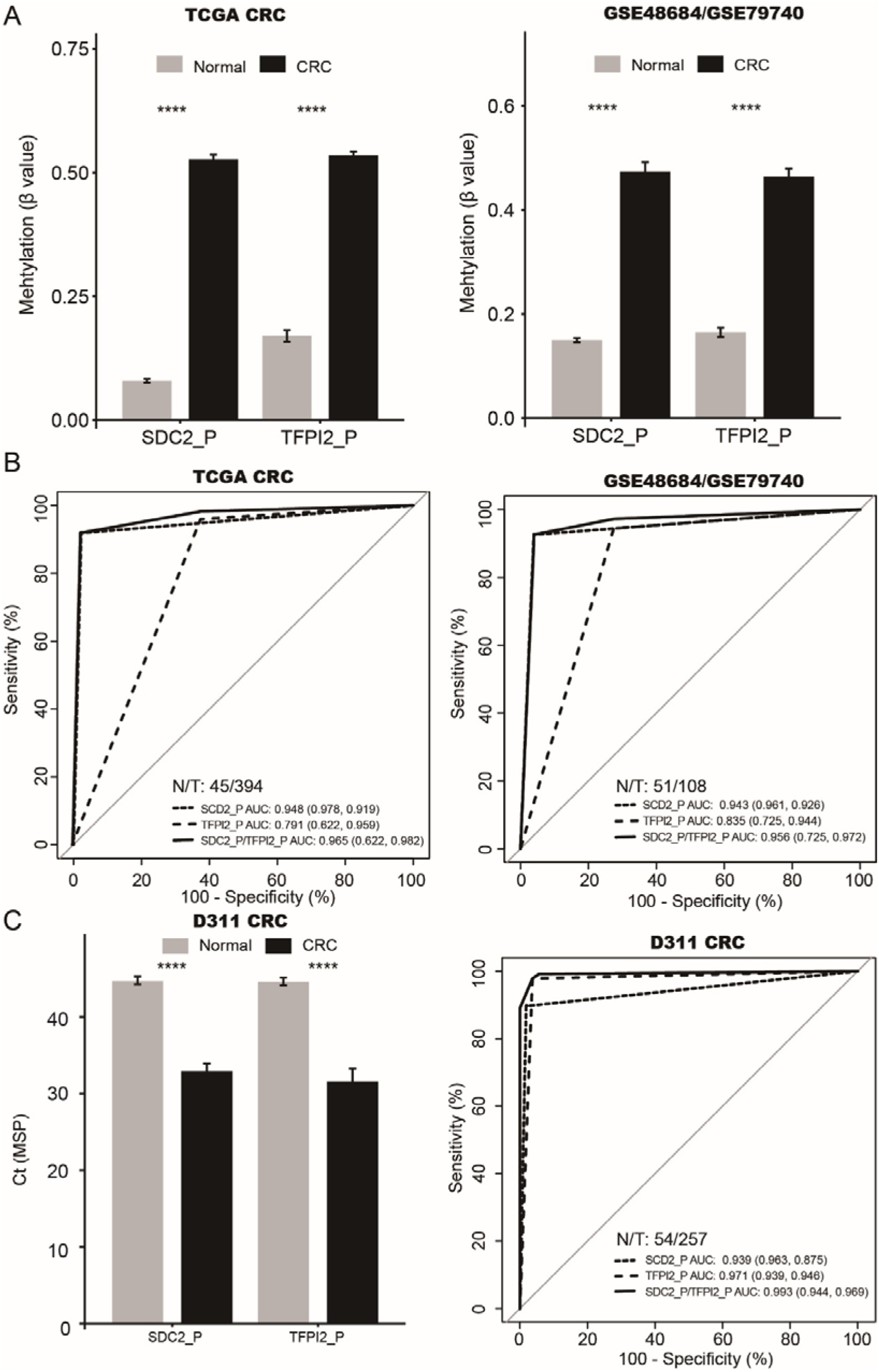
The performance of methylation levels of *SDC2* and *TFPI2* in distinguishing CRCs from normal controls_◦_ A: β values of SDC2 and *TFPI2* in CRCs and normal controls_◦_ B: ROC curves and area under the curve to evaluate the performance of SDC2_P and TFPI2_P for distinguishing CRC from normal samples. C: Ct values of SDC2 and *TFPI2* in normal and CRC samples in D311 dataset_◦_ Error bars represent mean β value±sd.

### HL group CRCs were more likely originated from left-side colon

We defined three methylator groups, HH (dual-positive), HL (single positive) and LL (dual-negative) according the promoter methylation status of *SDC2* and *TFPI2* (see methods). During the development of diagnostic biomarkers, CRCs divided to HL and LL groups are very important because of their effect on the sensitivity which might reflect a preference of the biomarkers for certain CRC subgroups. We first compared the three methylator groups with tumor location, and found that HL group CRCs were more frequently originated from left-side colon. A small amount of CRCs were from rectum, however very few were from right-side colon (**Figure 2A-C, Table 4**). This result indicated a potential impact of tumor locations on the early detection of CRCs by using *SDC2/TFPI2* dual-targets.

**Figure 2.**
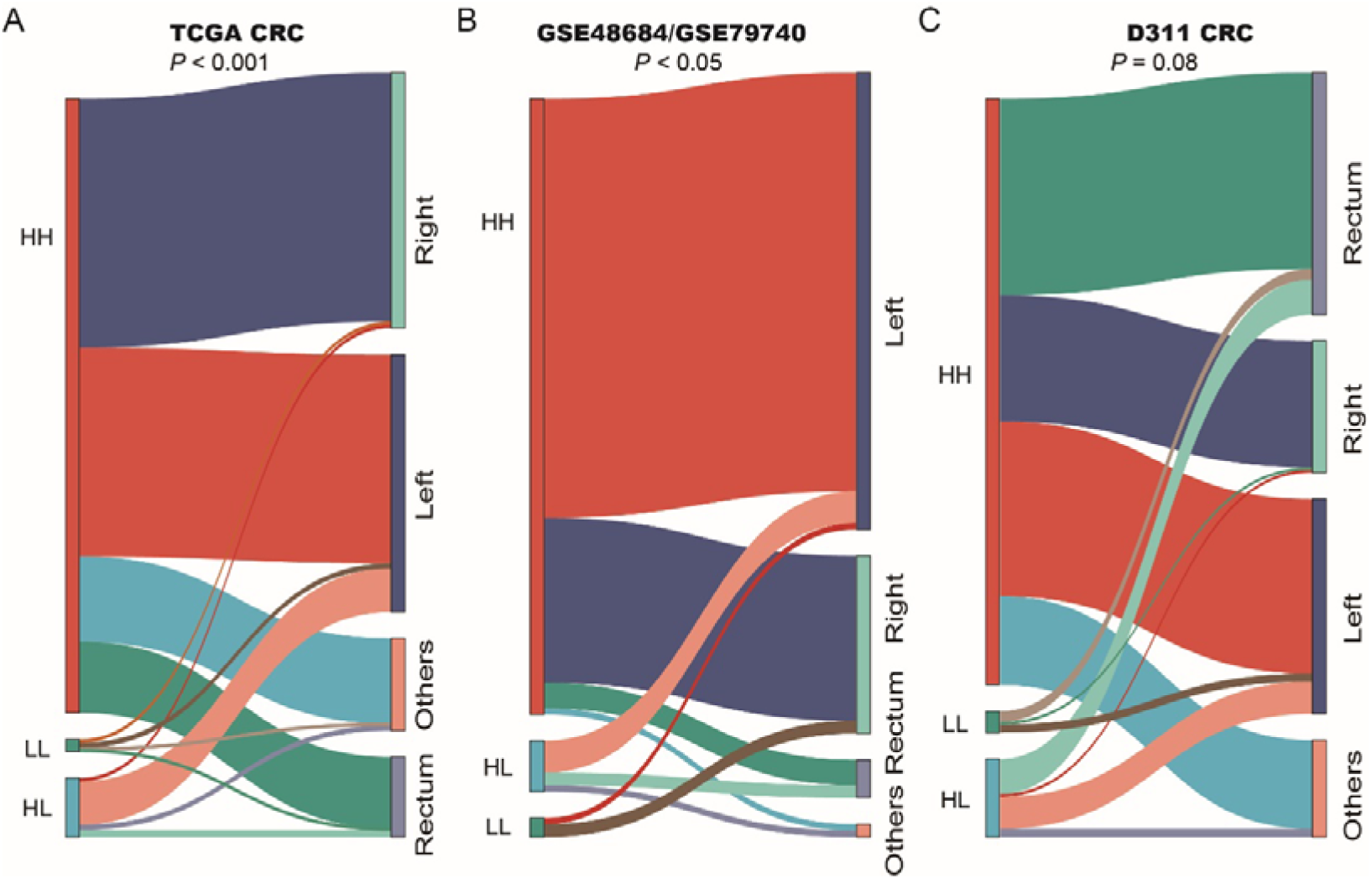
Sankey diagram showing the relationship of 3 methylator groups with tumor locations. Size of each rectangle and thickness of the connecting lines represents the number of samples on each group. Fisher exact test was used to calculate the *P* value.

**Table 4.** Samples in three methylator groups compared with tumor locations

| Tumor location | TCGA CRC |  |  | GSE48684/GSE79740 |  |  | D311 |  |  |
| --- | --- | --- | --- | --- | --- | --- | --- | --- | --- |
|  | HH | HL | LL | HH | HL | LL | HH | HL | LL |
| Right-side | 143 | 2 | 2 | 26 | 0 | 2 | 47 | 1 | 1 |
| Left-side | 120 | 25 | 3 | 66 | 5 | 1 | 65 | 12 | 3 |
| Rectum | 41 | 4 | 1 | 4 | 2 | 0 | 73 | 14 | 4 |
| Other | 49 | 3 | 1 | 1 | 1 | 0 | 34 | 3 | 0 |
| $\chi^2 P$ | 0.00059 | | | 0.015 | | | 0.081 | | |

### HH group CRCs presented more frequently genomic instability

Microsatellite instability and hypermutation have been regarded as important molecular characteristics of CRCs. Compared to other two groups, HH group CRCs presented the highest mutation load (**Figure 3A**, P < 0.05) in TCGA CRC dataset. By using the MANTIS score [46], which is used to evaluate the MSI status, we grouped the TCGA CRCs into MSI-H and MSS if this score is > 0.4. The β values of SDC2 and *TFPI2* showed a high concordance with MANTIS score in MSI-H group (**Figure 3B**, P < 0.001), which is consistent with the result that higher mutation load occurred in HH group. We studied the association of three methylator groups with the mutation of 5 typical CIMP-related genes including BRAF, PIK3CA, KRAS, TP53 and APC. All BRAF mutated CRCs were in HH group (**Figure 3C**, HH/HL/LL: 51/0/1, P = 0.018). We further compared the association between MSI-status and tumor locations with TCGA CRCs and our D3111 CRCs. The MSI-H CRCs were more preference in the right-side colon (**Figure 3C&D**, P < 0.001), which possibly elucidated potential causal factors that HL group CRCs were mainly in left-side colon and MSI-H CRCs were less likely in HL group.

**Figure 3.**
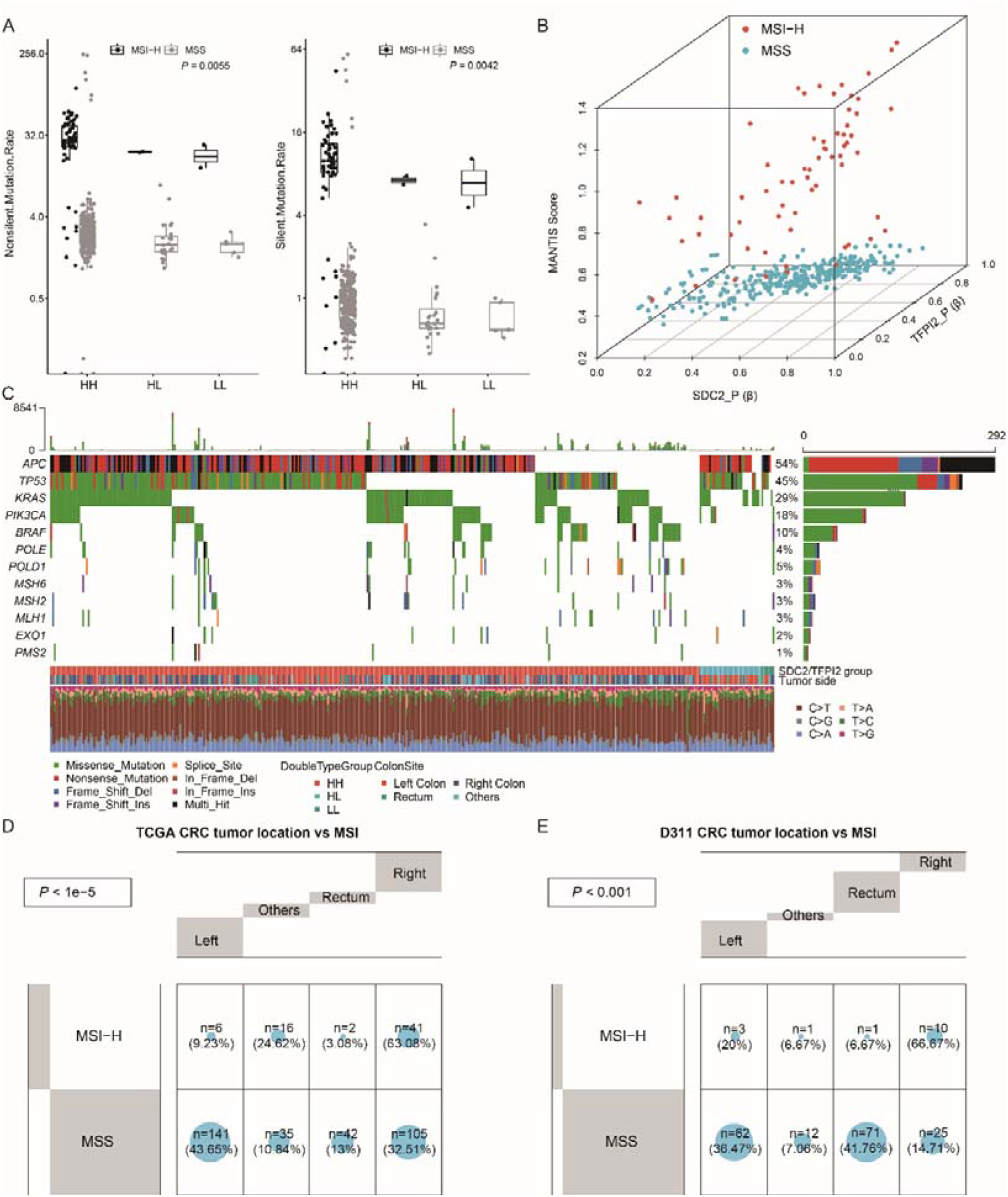
Genomic characteristics in three methylator groups. A: Nonsilent and silent mutation rates in HH, HL and LL groups. B: The correlation of MANTIS score with β values of *SDC2* and *TFPI2* in TCGA CRCs. C: Somatic mutation profile of five CIMP-related and MMR-related genes in HH, HL and LL groups. D&E: Comparison of tumor location with MSI in TCGA CRCs (D) and D311 CRCs (E), significant p value is calculated using fisher exact test.

### Older patients age was found in HH group CRCs

The age of patients is one of the risk factor for colon cancer and we found a significantly older age in HH group patients, while it was the youngest for LL group patients (**Figure 4A&B**, P < 0.05). Since the genomic DNA methylation is associated with patient age, we observed a positive correlation of the methylation levels of SDC2_P and TFPI2_P with patient age (**Figure 4C&D**), indicating that young patients might be more likely to be undetected.

**Figure 4.**
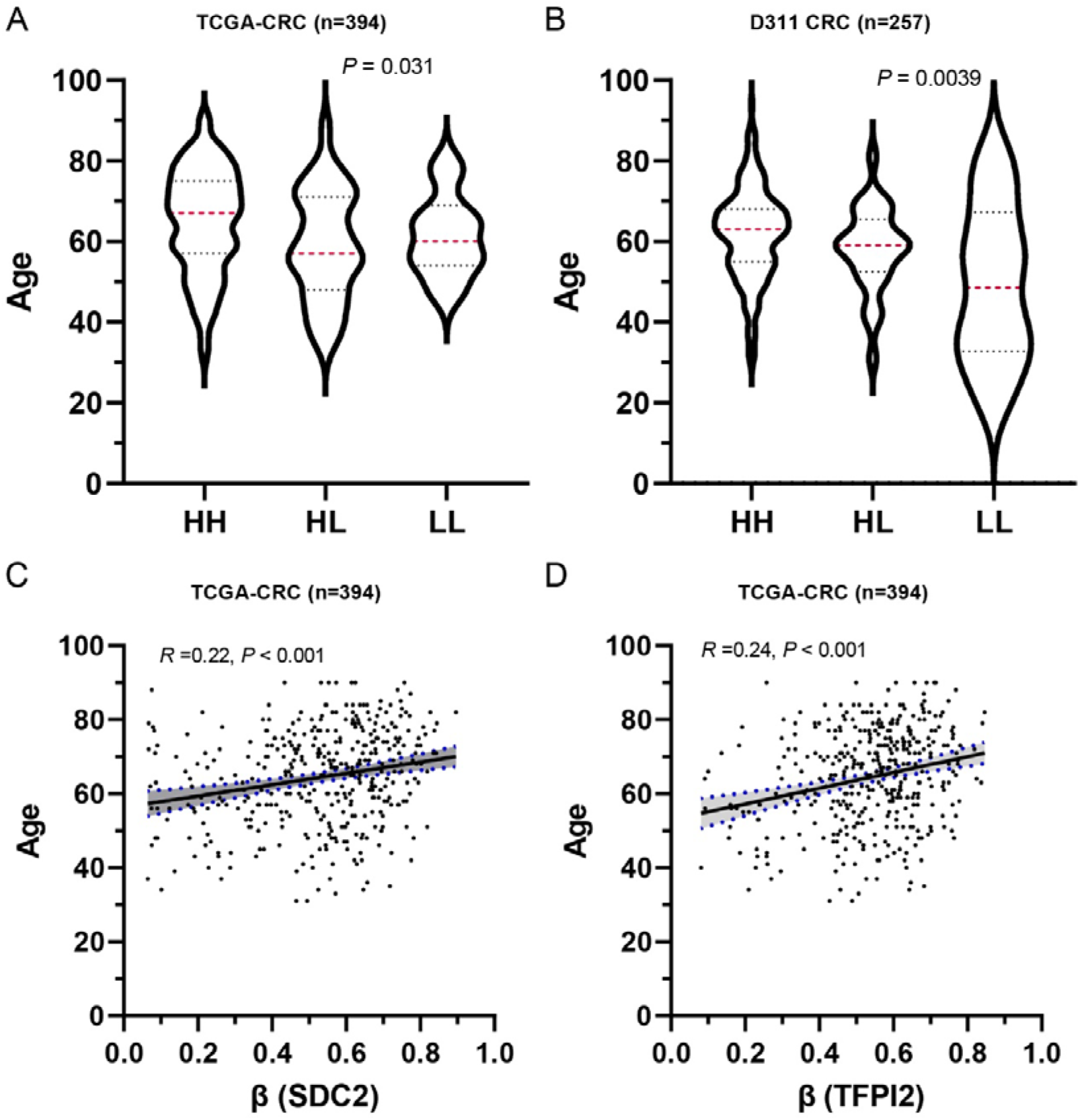
The association of patient age with methylation levels of *SDC2* and *TFPI2*. A&B: Patient ages in HH, HL and LL groups in TCGA CRCs (A) and D311 CRCs (B). The dashed red line indicate median age, while upper and lower dashed lines represent 75% quantile and 25% quantile. C&D: The correlation of patient age with β values of *SDC2* and *TFPI2* in TCGA CRCs. *P* value and correlation coefficient was calculated using pearsion’ method.

### LL group CRCs might be related to alterations of ECM and cell migration biological processes

We performed differential expression analysis by using the gene expression profile of TCGA CRCs to identify group-specific DEGs. A total of 37 HH specific, 84 HL specific and 22 LL specific DEGs were identified according to their average expression values on the three groups (**Figure 5A**). Functional enrichment analysis implied that HH specific DEGs were mainly related to the regulation of transcription and other processes (**Figure 5B**), while LL specific DEGs are enriched in the biological processes of extracellular matrix interaction (ECM) and cell migration (**Figure 5C**). These results might elucidate potential alterations in the biological processes of ECM and cell migration that lead to different characteristics of these three groups.

**Figure 5.**
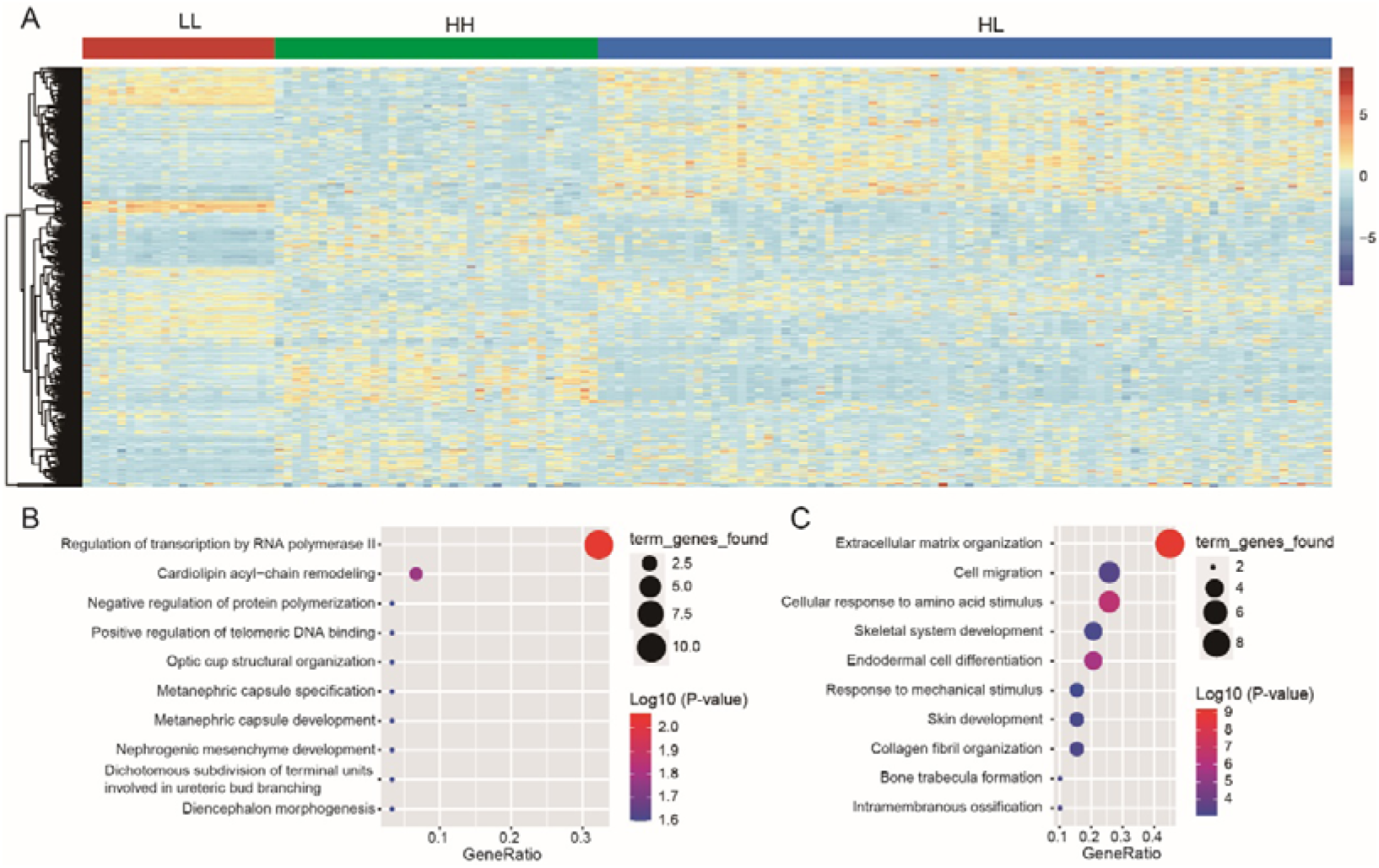
Identification of group specific DEGs. A: Expression heat map for the HH-, HL- and LL-specific DEGs. B: GO functional enrichment analyses for HH-specific DEGs. C: GO functional enrichment analyses for LL-specific DEGs. The log10 (P-value) of each term is colored according to the legend.

## Discussion

Quantifying aberrant methylated genes was useful and feasible method for the early detection of CRCs. However many of these biomarkers have the limitation of low sensitivity such as the undetected false negative samples [24,47]. In this study, we defined three CRC methylator groups, dual-positive (HH), single positive (HL) and dual-negative (LL) based on the methylation status of *SDC2* and *TFPI2* and then evaluated their characteristics of genomic instability, mutation load, patient age, and biological processes.

The performance of *SDC2* and *TFPI2* for discriminating CRCs from normal controls was first evaluated in three independent datasets and our preliminary study revealed a relatively higher specificity and lower sensitivity for detection of *SDC2* methylation than *TFPI2*. This might be attributed to the higher background methylation level in the promoter of *TFPI2*. The detected sensitivity was improved when combing the dual-targets compared to single target, indicating a complemental performance of SDC2 and *TFPI2* in early detection of CRCs. Previous studies have demonstrated that multi-target outperformed single target [48,49], which was also found in this study.

In clinical practice, the single-positive and dual-negative CRCs detected by dual-target will cut down the sensitivity of these biomarkers- Our findings indicated that single-positive CRCs more likely tend to originate from left-side colon. Several studies have demonstrated that left-side colon presents lower degrees of methylation, while right-side colon shows high degrees of methylation which was called the CpG island methylator phenotype, or CIMP [50]. This might explain why single-positive CRCs appear more frequently in left-side colon. Additionally, we found a positive correlation of the methylation levels of *SDC2* and *TFPI2* with MSI scroe in MSI-H CRCs, as well as lower mutation load and rare BRFA mutations in HL group CRCs. These results, on the other hand, confirmed that the molecular events, such as epigenetic instability, aberrant DNA mutation and MSI, are coupled with each other [25]. Gene expression analysis identified methylator group-specific DEGs and functional annotation of LL-specific DEGs was suggested to focus on the biological process of ECM-receptor interaction implying a potential alteration of molecular pathway in LL group CRCs. Interestingly, many studies have showed very important roles of SDC2 wan *TFPI2* in the interaction of extracellular matrix with cell plasma [15,51]. These findings revealed the possible impact of ECM process on the performance of *SDC2/TFPI2* dual-target in detecting CRCs.

Colorectal cancer is a disease with high heterogeneity. CRCs are often classified to proximal (right side) and distal (left-side) according to their anatomical locations. This classification is reasonable because of their distinctive embryonic derivation, which is the midgut and the hindgut for the proximal and distal colon, respectively [27,31]. It might be related to the differences in DNA methylation between left- and right-side colon, with potential impact on the detection of CRCs based on methylation status. In summary, the current study demonstrated the possible association of CIMP phenotype, tumor location and MSI with the dual-target in CRC early diagnosis. Our observations also suggested that it should be considered during the development of new methylation based biomarkers for CRC detection in these respective sides.

## Acknowledgments

This study was supported by Medical Science and Technology Research Plan Joint Construction Project of Henan Province (Granted No. 2018020121). We declare that submitted manuscript and other materials are not under consideration for publication elsewhere. All authors listed have read the complete manuscript and have approved submission of the paper.

## Conflict of Interest

All authors declare no conflict of interest.

## Data Availability Statement

The TCGA CRC 450k data and GEO datasets are publicly available online. The D311 data used and analyzed in study are available from the corresponding author on reasonable request.

## Author Contributions

Study design: HJ Y.

Data collection and analysis: RX L, YT Z, KK W and TT L.

Manuscript writing: RX L, YT Z, K H and KK W.

Manuscript and results revising: HJ Y and XP L.

## Notes

### Competing Interest Statement

The authors have declared no competing interest.

